## Supplementary Information for "Ais: streamlining segmentation of cryo-electron tomography datasets"

---

### Supplementary Figures

Supplementary Figures 1 - 5 show the Ais interface for the various steps in the segmentation workflow: annotation (1), training models (2), testing models (3), exporting volumes (4), and inspecting the results (5). To illustrate the duration of processing, the time spent preparing each step is noted alongside the figure title. For optimal results it is best to spend more time on each step - the time spent for this example is meant to give an indication of how quickly one can achieve a first result.

#### Figure S1 - Annotation & boxing interface (3 minutes)

The image shows an example of membranes (red) being segmented and positive (red boxes) and negative (yellow boxes) training boxes being placed. Neural networks in Ais operate on square input images (here 64 x 64 pixels), and the training data thus consists of square pairs of images, with the grayscale data in a placed box as the training input and the user-drawn annotations in that same box as the training output.

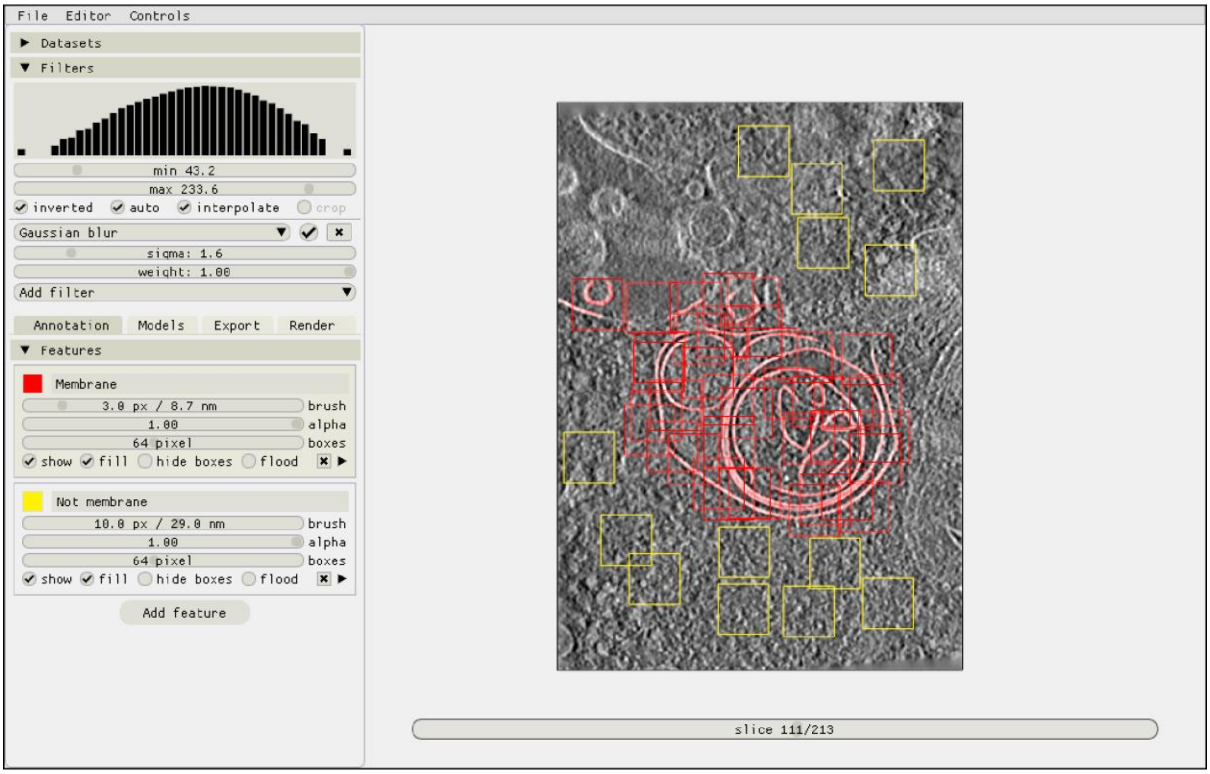

**Figure S2 - Exporting training datasets & training models (2 minutes)**

After annotation and boxing, training datasets can be exported by selecting which annotations to use as positives and which as negatives. In this example, the feature 'Membrane' is included as a positive feature, meaning input grayscale and output annotation are sampled. The feature 'not membrane' is included as a negative, meaning input grayscale is sampled and the corresponding output annotations are all zero. Including boxes for the 'membrane' feature that do not show membrane in the grayscale image and also are not annotated has the same effect as using separate positive and negative classes.

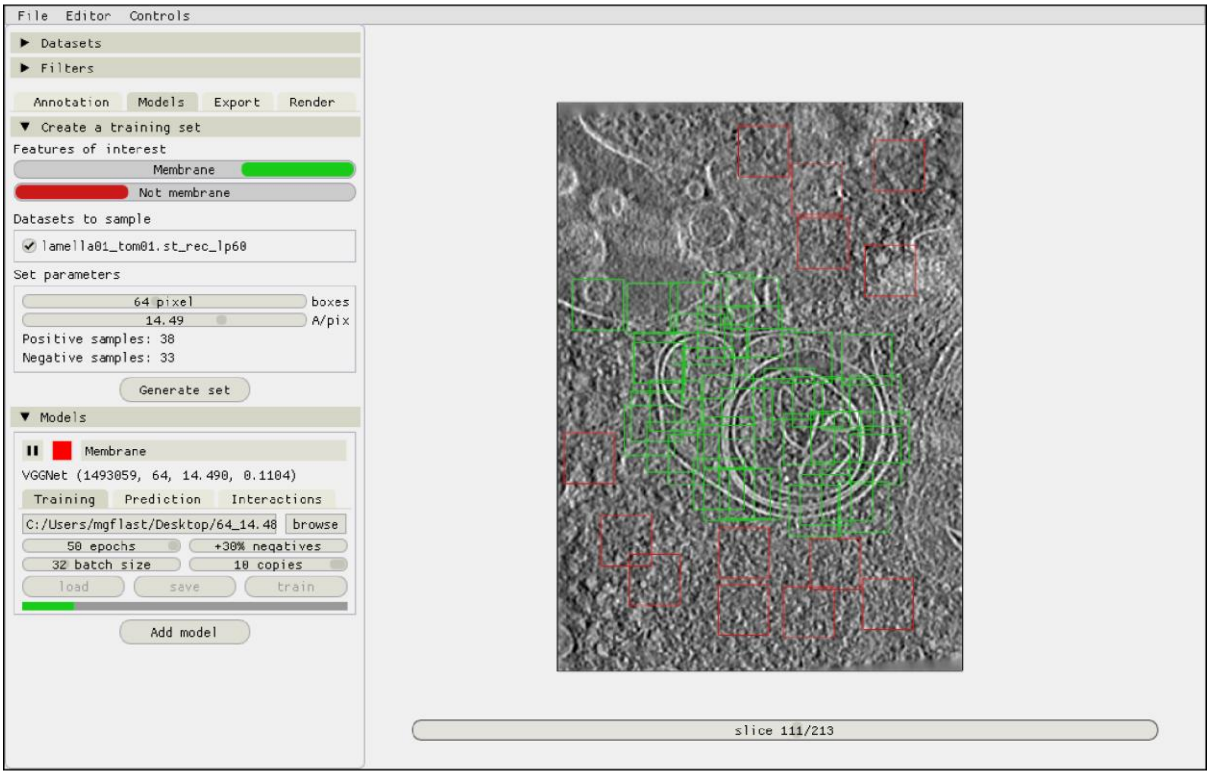

### Figure S3 - Testing a model (1 minute)

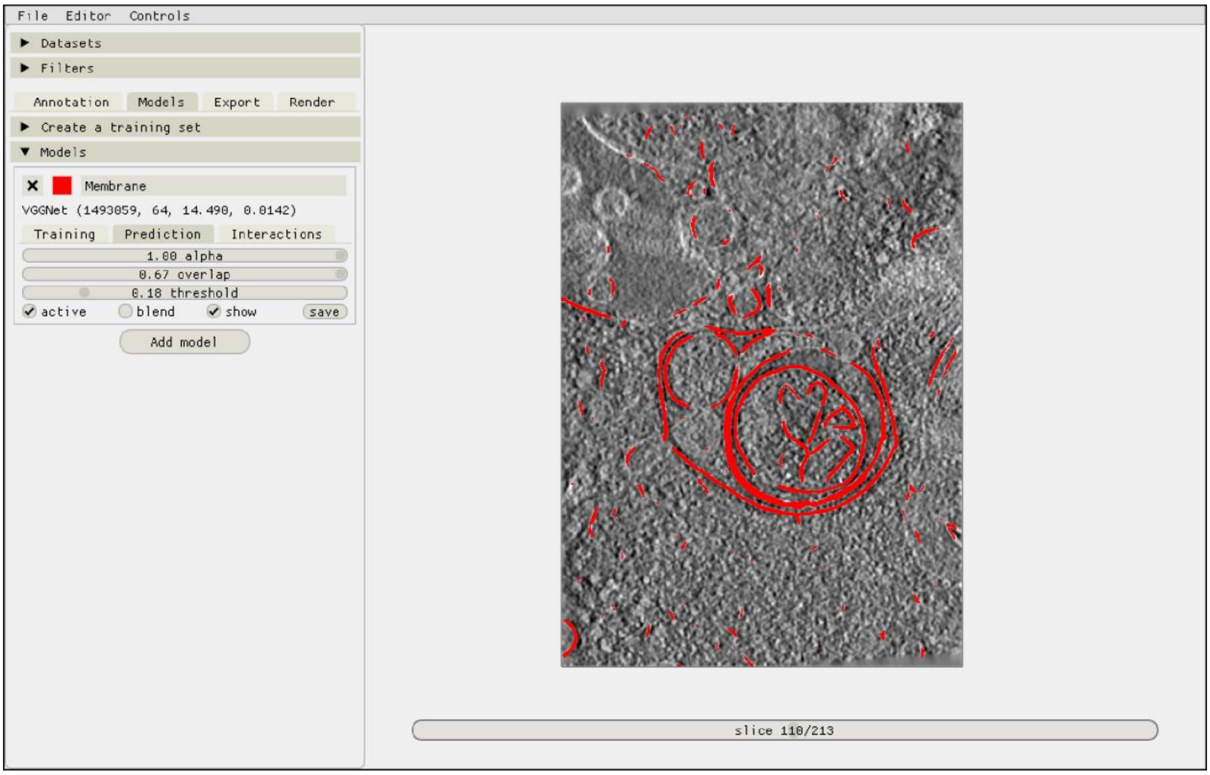

### Figure S4 - Exporting a volume (1 minute)

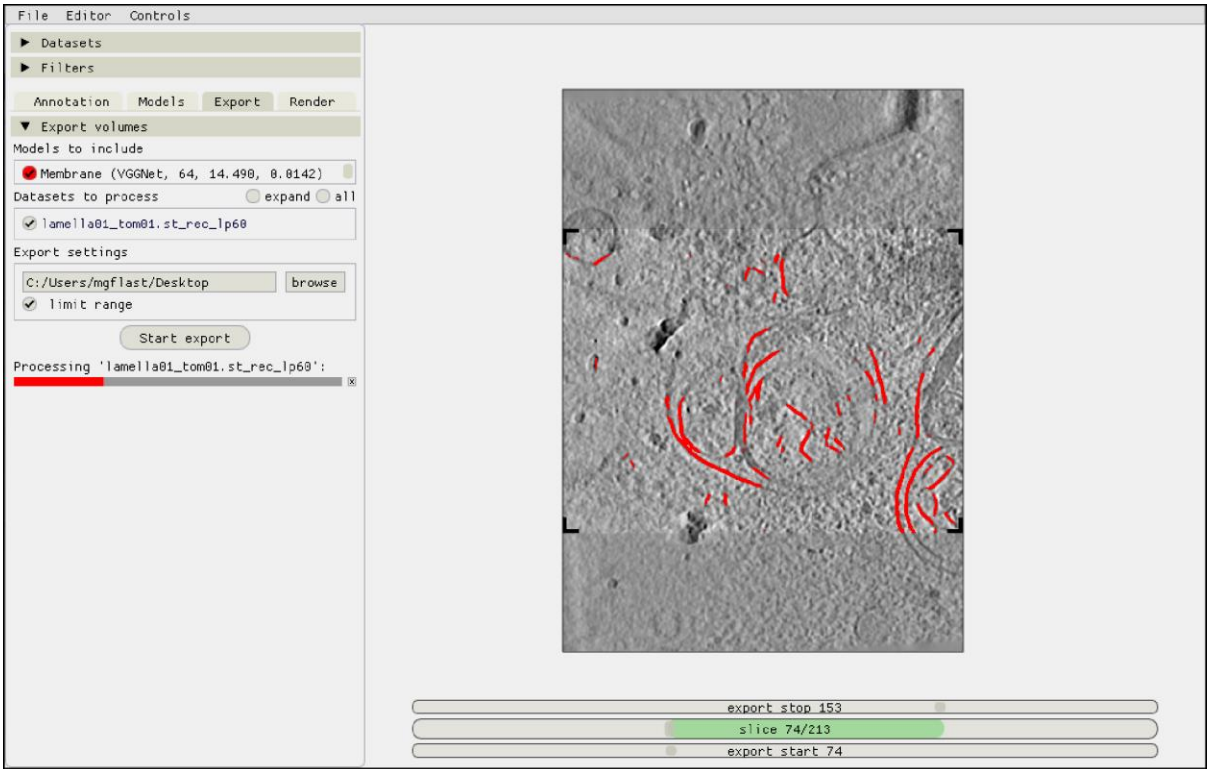

### Figure S5 - Inspecting the result (< 1 minute)

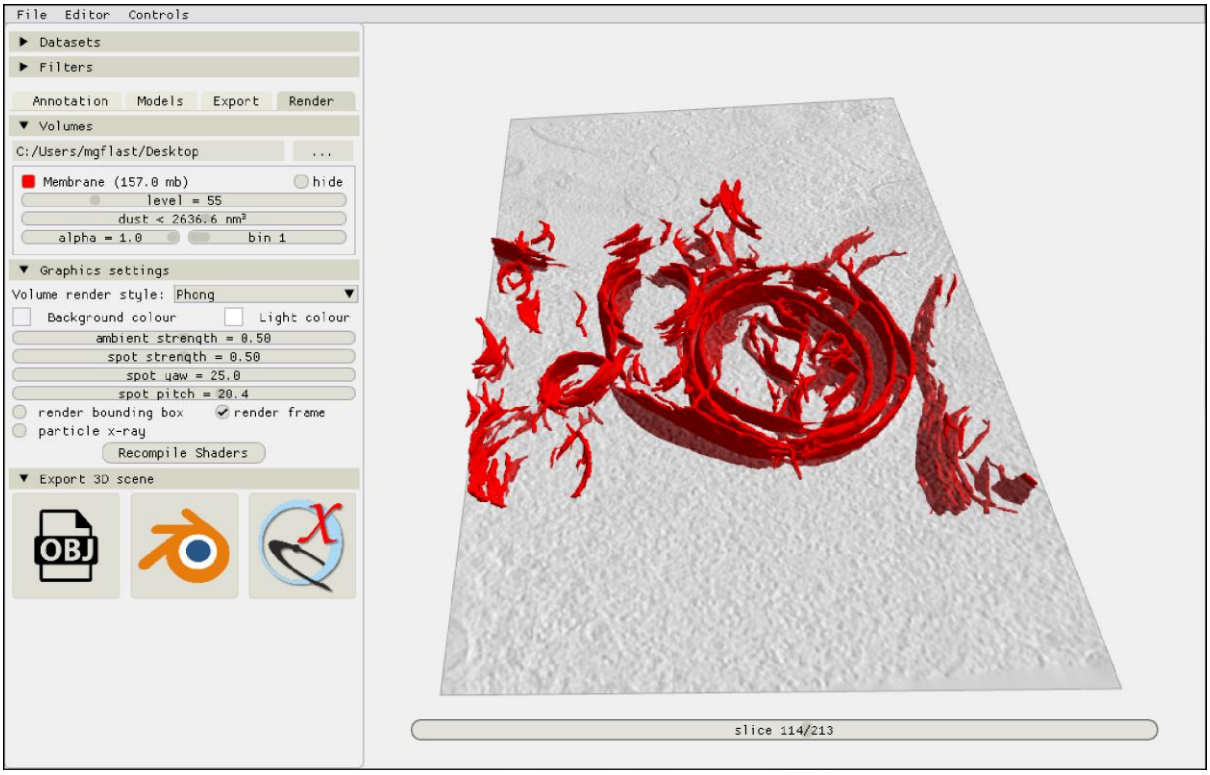

**Figure S6 – Comparison of manual annotations and UNet, Pix2pix, and improved Pix2pix antibody platform segmentations for the data in Figure 2.**

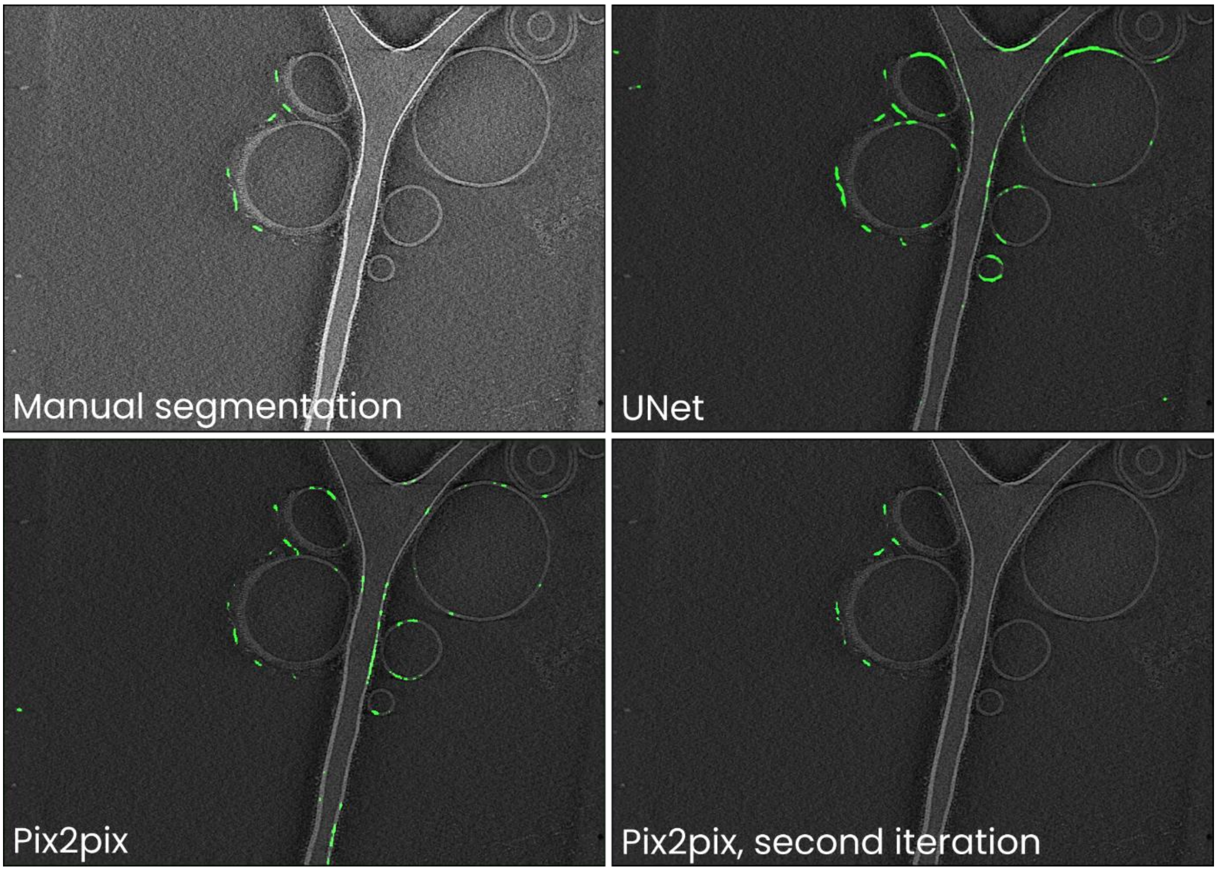

Training neural networks can often be an iterative process: an initial training dataset is compiled, a model trained, and the resulting segmentations are likely to contain readily identifiable false negatives (i.e. parts of antibody platforms not annotated by a model) and positives (e.g. membranes segmented as antibody platforms). To improve the model, it is then useful to perform a second iteration of annotating boxes for use in training and then training the model anew. In Ais, this is facilitated by being able to switch back and forth between the 'annotation' and 'models' tabs (**Figs. S1-S3**), and by being able to quickly test models on and annotate new boxes in many different datasets.

In the example above, we initially trained Pix2pix on the same training dataset as used for the other networks discussed in Figure 2 and Table 1 in the main text. Although the loss of the UNet network was the lowest, we found that in comparison to most other networks the output of the Pix2pix model contained fewer false positive predictions (e.g. compare the segmentations of the liposome membranes). Since Pix2pix is a large model, it is likely to benfit the most from further training on an expanded training dataset. We thus included additional samples in the training dataset and trained a new instance of a Pix2pix model for 50 epochs. The resulting model produced segmentations that were much closer to the manual annotations.

Further information and tips for preparing useful models are also discussed in the video tutorials, available at: [youtube.com/@scNodes](https://youtube.com/@scNodes)

**Figure S7 – A visual (2D) representation of the processing steps employed in the automated picking process**

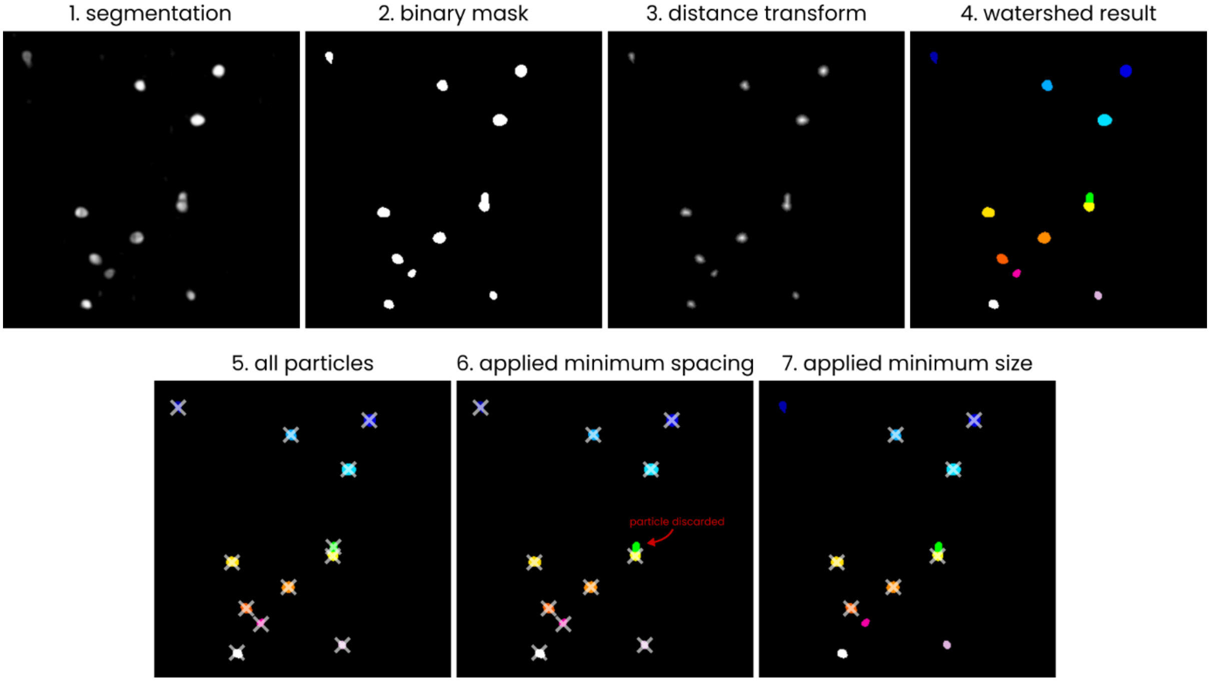

The steps employed in the process of converting a segmented .mrc volume to a list of particle coordinates. The example is shown in 2D; the actual picking process is performed in 3D. 1) The input segmentation (values 0 – 255), 2) a binary mask, generated by thresholding the input image at a value of 127, 3) distance transform of the binary mask, 4) groups of pixels, labelled using a watershed algorithm (skimage.segmentation.watershed) with local maxima in the distance transform output used as sources, 5) centroid coordinates (marked by crosses) of the pixel groups, 6) example of removing particles by applying a minimum spacing, 7) example of removing particles by specifying a minimum particle size.

### Figure S8 - Inspecting the autopicking results in the Ais renderer

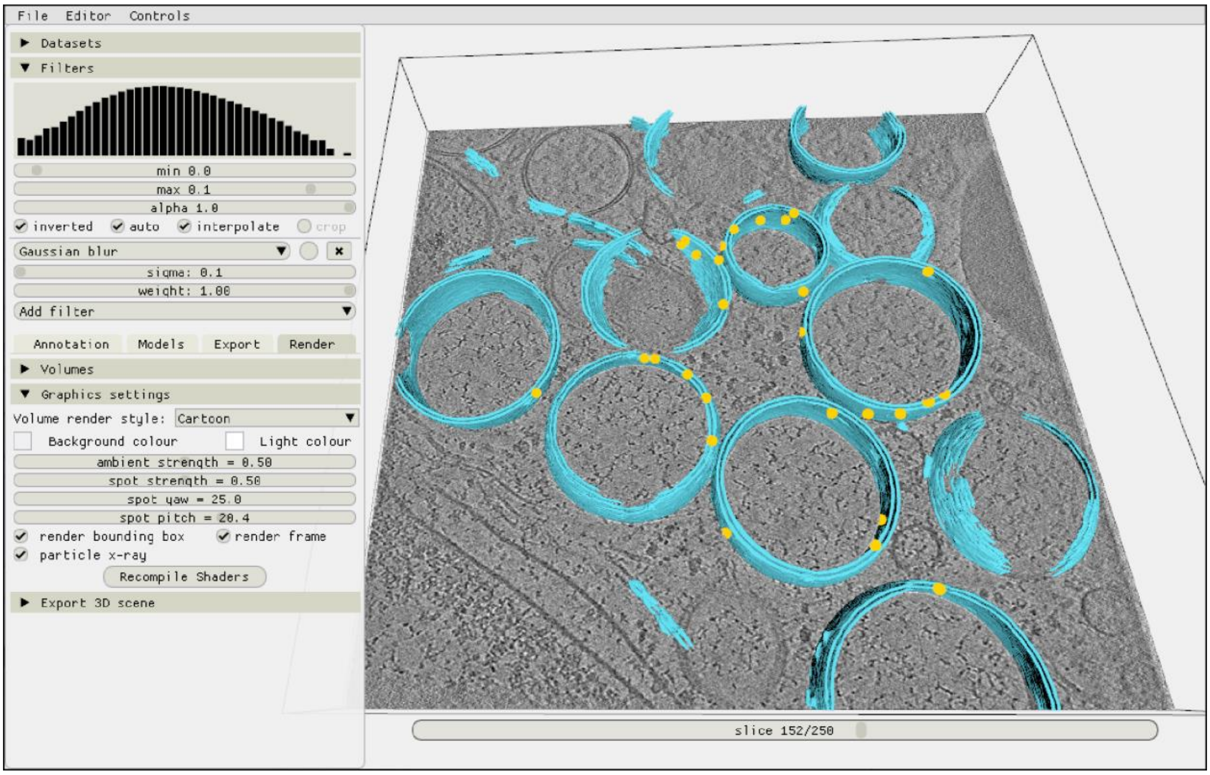

After picking particles, the resulting localizations are immediately available for inspection in the renderer. In this example, yellow markers indicate the coordinates of molecular pores, picked in a tomogram from the dataset by Wolff et al. (see main text Figure 5).

**Figure S9 - Comparison of the automatically picked C1-IgG3 complex reconstruction versus the original reconstruction in Abendstein et al. 2023.**

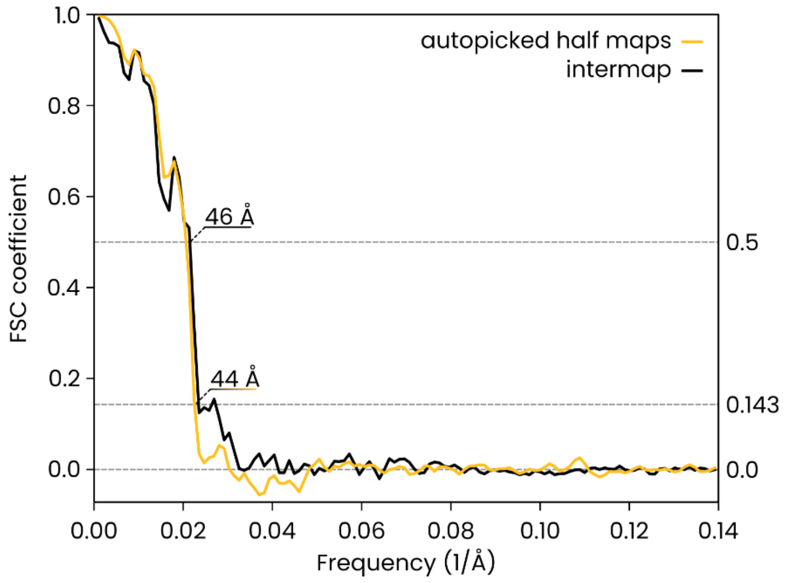

The FSC curve for the two half maps of the automatically-picked C1-IgG3 complex reconstruction gives a resolution of 44 Å at the FSC =0.143 (yellow, 'autopicked half maps'). In the original publication (Abendstein et al. (2023), Nat. Comm., doi.org/10.1038/s41467-023-39788-5) various resolutions are reported for different maps that were obtained by focused refinement on different parts of the complex. In the current report, we did not apply further refinement steps beyond the initial even/odd reconstructions on the entire complex that the above FSC curve is based on. The value of 44 Å must thus be compared to the corresponding initial reconstruction in the original report (in their Supplementary Fig. 10, panel c, map #1) which was reported at FSC = 0.143 with the same resolution of 44 Å.

A comparison between the full maps (black FSC curve, 'intermap') at FSC coefficient 0.5 gives a value of 46 Å for the resolution, thus indicating that the two independent reconstructions did indeed converge on the same structure.

**Figure S10 - Individual images of the ten features segmented in the coronavirus infected mammalian cells dataset (Wolff et al., 2020, Science)**

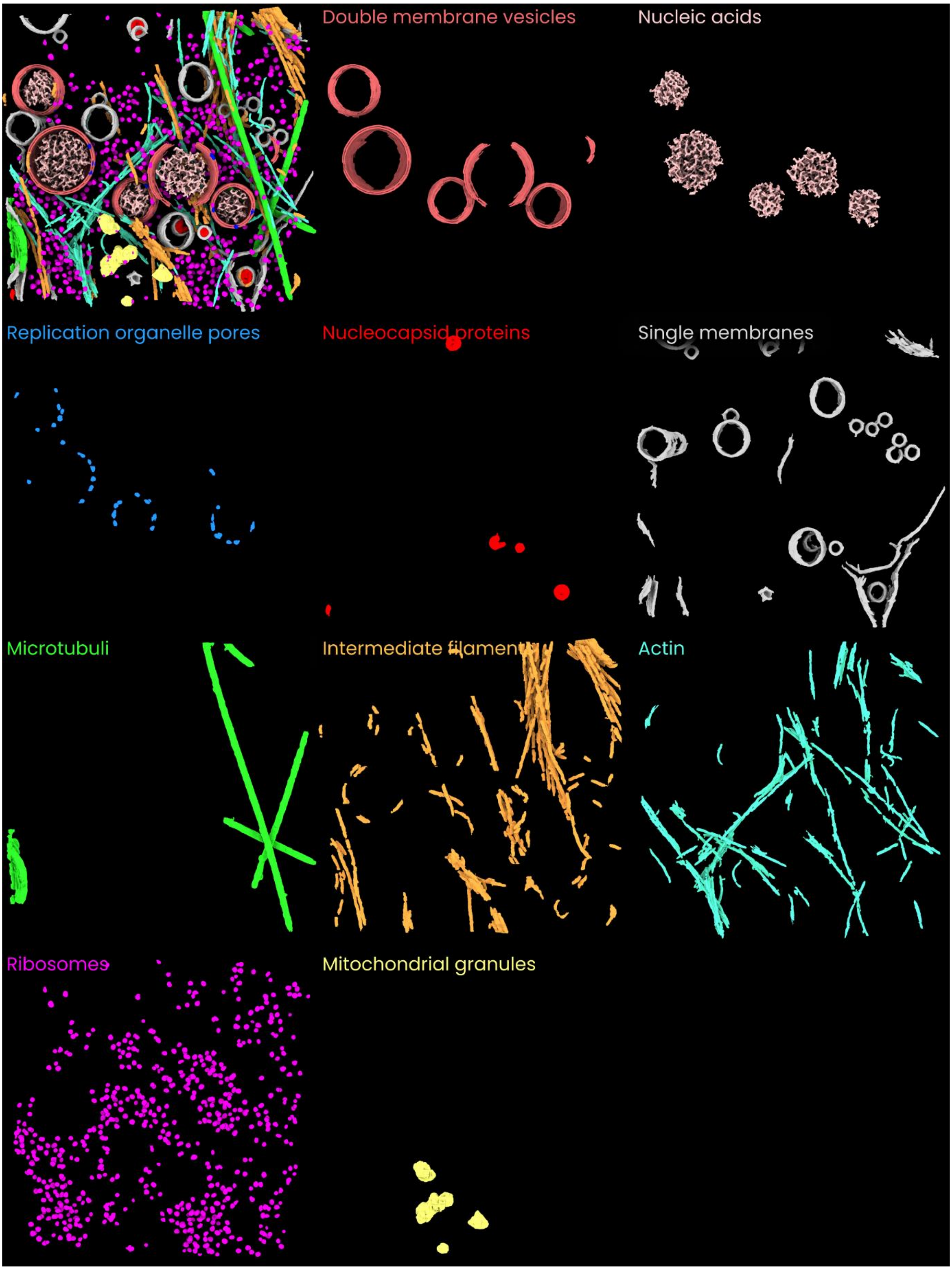

**Figure S11 - A collage of automatically picked molecular pores in the double membrane vesicles**

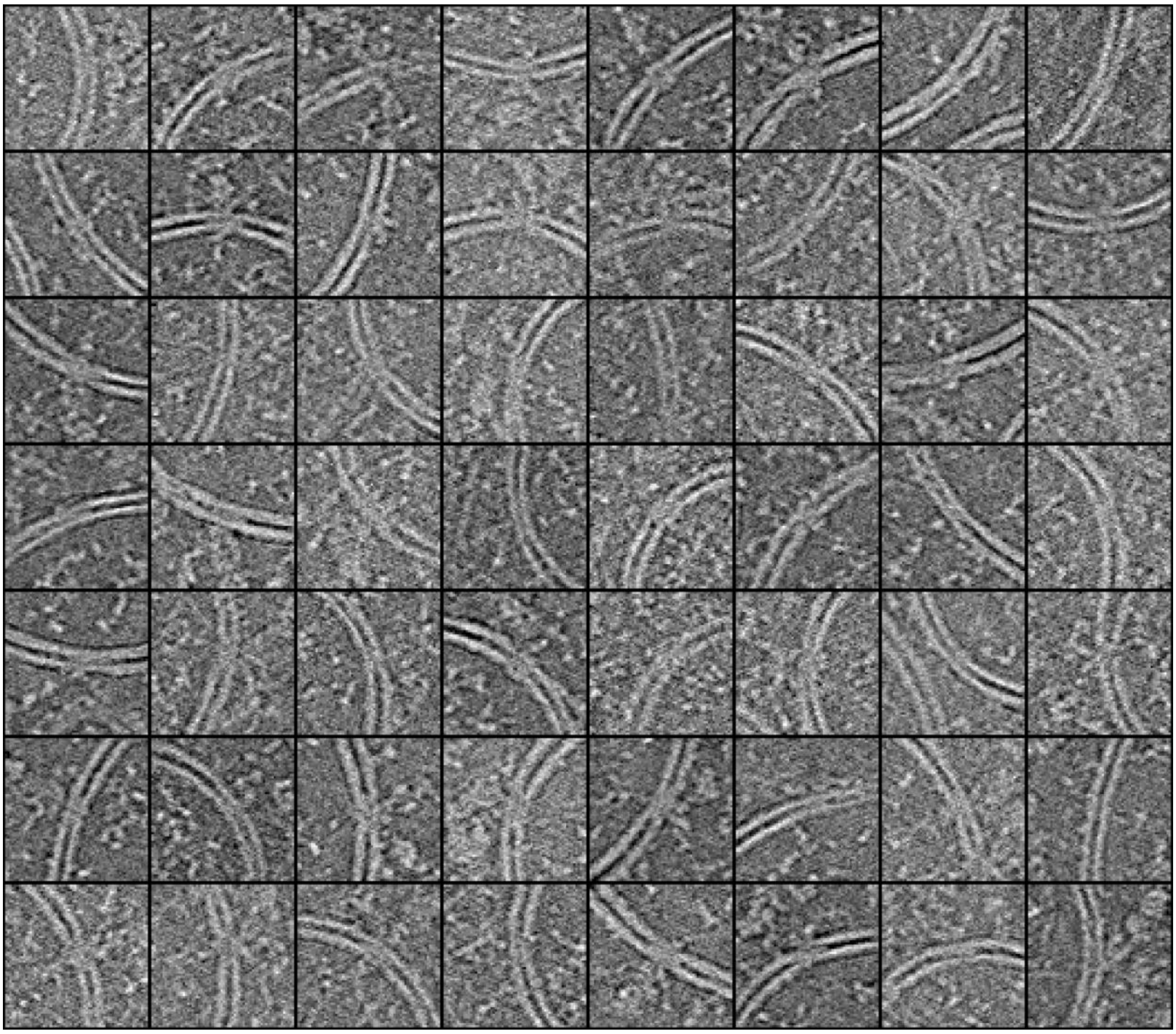

As with the C1-IgG3 complexes and IgG3 platforms (main Figures 3 and 4), the segmentations of the molecular pores in the coronavirus replication organelle (main Figure 5) can be used to automatically pick particles. In the above image, tomographic slices are shown that were cropped at the location of 56 particles (picked in two tomograms). Although it can be difficult to tell at the single particle level whether an image contains the exact structure of interest, the images are all centered on sections of double membrane vesicles and many show additional density between the membranes as well as at the outside (convex side) of the vesicle, characteristic of the molecular pore.

The accuracy of picking and the quality of the resulting particle datasets are dependent on the quality of the original segmentations and on the parameters used in picking. Converting a 3D grayscale volume to a list of coordinates requires a number of parameters to be specified (in Ais, but also in many other programs). First, a threshold value is used to convert from grayscale to a binary volume. Contiguous regions of nonzero voxels in the volumes are then considered 'particles', and various metrics can be used to determine whether particles are included in the final list of coordinates. Example are the particle volume, integrated prediction value for the voxels included in the particle's volume, or a minimum particle spacing. In Ais, users can specify the first and the latter (we experimented with using the second metric as well, but found it very similar but less intuitive than the metric of particle volume, so we did not include it). The threshold values chosen for these metrics and for the conversion of the grayscale volume to a binary one thus define what is ‘dust’. In practice, choosing these values entails a trade-off between the number of particles, and the ‘quality’ of the selected set of particles. Low thresholds yield a large number of particles, which can introduce the need to perform classification during reconstruction by subtomogram averaging. High thresholds yield fewer particles, but the resulting selection may be more uniform and their images of higher quality. In practice, it is advisable to test different thresholds and to inspect the resulting particle datasets, in order to ensure that i) good particles are not being excluded, and ii) large numbers of bad particles (e.g. densities that correspond to structures other than the structure of interest) are not being included. As with manually picked particle sets, classification is likely required to curate the selection of particles.

### Supplementary Note 1: Python file format for Ais models

#### 1.1 Adding a Keras model

Most models in the standard Ais library are Keras models (tensorflow.keras). Adding an extra keras model with a new architecture is relatively straightforward and can be achieved by adding a .py file to Ais/models directory. The .py file requires three components: a title for the model, a boolean that specifies whether the model should be available in the software, and a function 'create' that returns a keras model. The implementation of the VGGNet model (vggnet.py) is copied below as an example.

```python
from tensorflow.keras.models import Model
from tensorflow.keras.layers import Input, Conv2D, MaxPooling2D, Conv2DTranspose
from tensorflow.keras.optimizers import Adam

title = "VGGNet"
include = True

def create(input_shape):
    inputs = Input(input_shape)

    # Block 1
    conv1 = Conv2D(64, (3, 3), activation='relu', padding='same')(inputs)
    conv2 = Conv2D(64, (3, 3), activation='relu', padding='same')(conv1)
    pool1 = MaxPooling2D(pool_size=(2, 2))(conv2)

    # Block 2
    conv3 = Conv2D(128, (3, 3), activation='relu', padding='same')(pool1)
    conv4 = Conv2D(128, (3, 3), activation='relu', padding='same')(conv3)
    pool2 = MaxPooling2D(pool_size=(2, 2))(conv4)

    # Block 3
    conv5 = Conv2D(256, (3, 3), activation='relu', padding='same')(pool2)
    conv6 = Conv2D(256, (3, 3), activation='relu', padding='same')(conv5)
    pool3 = MaxPooling2D(pool_size=(2, 2))(conv6)

    # Upsampling and Decoding
    up1 = Conv2DTranspose(128, (2, 2), strides=(2, 2), padding='same')(pool3)
    conv7 = Conv2D(128, (3, 3), activation='relu', padding='same')(up1)

    up2 = Conv2DTranspose(64, (2, 2), strides=(2, 2), padding='same')(conv7)
    conv8 = Conv2D(64, (3, 3), activation='relu', padding='same')(up2)

    up3 = Conv2DTranspose(1, (2, 2), strides=(2, 2), padding='same')(conv8)
    output = Conv2D(1, (1, 1), activation='sigmoid')(up3)

    # create the model
    model = Model(inputs=[inputs], outputs=[output])
    model.compile(optimizer=Adam(), loss='binary_crossentropy')

    return model
```

#### 1.2 Adding a custom model

Adding a non-Keras model is also possible but requires a little bit of extra work. Only a small number of methods of the Keras model object type are directly accessed by Ais. These are: count_params, fit, predict, save, and load. Adding a custom model thus requires adding a .py file to the *Ais/models* that contains four components: a title, a boolean that specifies whether the model is available in the software, and a function 'create' that returns model object (these are as before, with adding a keras model), and additionally a definition of a class that implements the required methods. The return types of these methods should be the same as those returned by the corresponding Keras methods. The content of the *model_template.py* template file is copied below as an example.

```
title = "Template_model"
include = False

def create(input_shape):
    return TemplateModel(input_shape)

class TemplateModel:
    def __init__(self, input_shape):
        self.img_shape = input_shape
        self.generator, self.discriminator = self.compile_custom_model()

    def compile_custom_model(self):
        # e.g.: compile generator, compile discriminator, return.
        return 0, 0

    def count_params(self):
        # e.g. return self.generator.count_params()
        # for the default models, the number of parameters that is returned is the
amount that are involved in processing, not in training. So for e.g. a GAN, the
discriminator params are not included.
        return 0

    def fit(self, train_x, train_y, epochs, batch_size=1, shuffle=True,
callbacks=[]):
        for c in callbacks:
            c.params['epochs'] = epochs

        # fit model, e.g.:
        for e in range(epochs):
            for i in range(len(train_x) // batch_size):
                # fit batch
                pass

                logs = {'loss': 0.0}
                for c in callbacks:
                    c.on_batch_end(i, logs)

    def predict(self, images):
        # e.g.: return self.generator.predict(images)
        return None

    def save(self, path):
        pass

    def load(self, path):
        pass
```

A more concrete example of the implementation of a custom model can be found in Ais/models/pix2pix.py. The pix2pix model is implemented in Keras, but since it internally requires the use of two separate Keras model objects (the generator and the discriminator), implementing it in Ais was a matter of wrapping the pix2pix models in a custom class. See: https://github.com/bionanopatterning/Ais/blob/master/Ais/models/pix2pix.py

#### 1.3 Importing and exporting models

**Export**

After training, Ais models can be saved for future re-use or for uploading to the repository aiscryoet.org. The resulting savefiles, with extension *.scnm*, are uncompressed .tar archives that contain an number of required files:

- A .h5 file that fully describes the CNN architecture and weights. *The .h5 file is generated using the standard keras procedure model.save.*
- A .json metadata file, which contains information on the name and processing parameters used for the model.

If a tomogram was open in Ais at the moment of saving the model, two additional files are also included:

- A .tiff file, containing a single slice from that tomogram, saved so that the model performance can be validated prior to releasing it on the repository.
- A .png file, an image (downsized 512 × 512 pixels) depicting the validation slice with a segmentation overlaid on top, which is used as the thumbnail on the model repository.

Of these last two files, only the .png file is publically available on the repository. The image that will be publically visible is also displayed on the aiscryoet.org/upload page, prior to actually uploading the model.

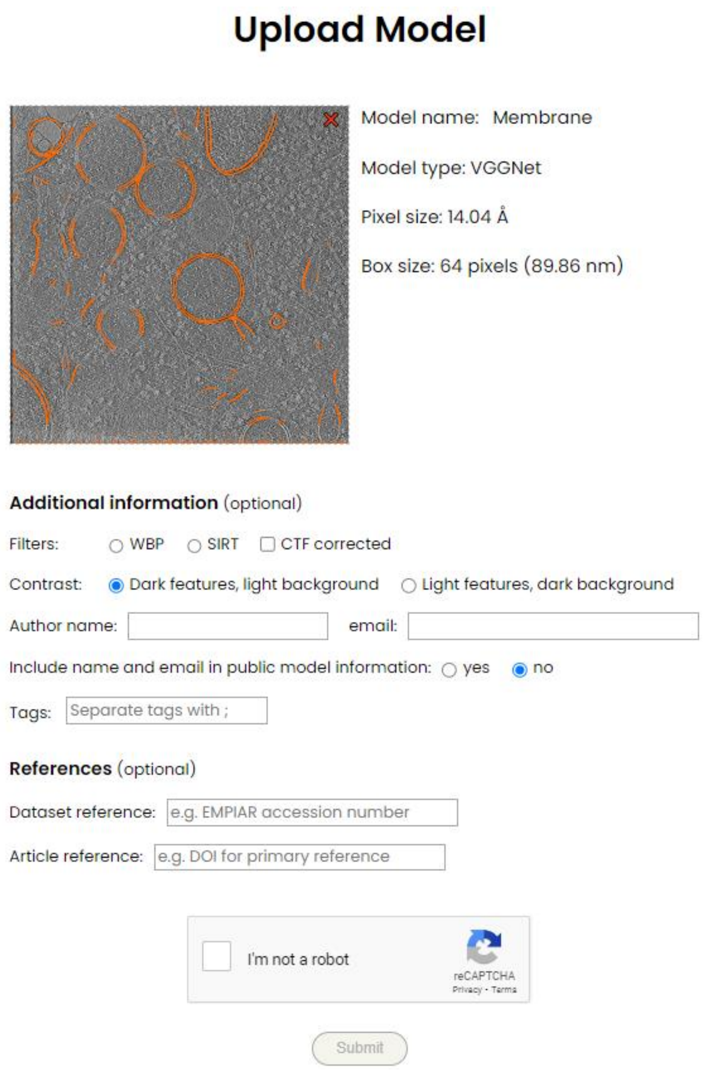

Figure S12 – Uploading a model to the model repository via aiscryoet.org/upload.

**Import**

A .scnm tar archive that contains correctly formatted .h5 and .json files can be loaded into Ais. Models generated elsewhere can also be imported, if these two files are combined into a .tar archive and renamed .scnm.

*The .h5 file is read using the standard keras load_model procedure.*

The following content is expected in the .json:

```
{
    "title": "Membrane",
    "colour": [1.0, 0.407843142747879O3, 0.0],
    "apix": 14.04,
    "compiled": true,
    "box_size": 64,
    "model_enum": 5,
    "epochs": 50,
    "batch_size": 64,
```

```
    "active": true,
    "blend": false,
    "show": true,
    "alpha": 0.75,
    "threshold": 0.5,
    "overlap": 0.20000000298023224,
    "active_tab": 0,
    "n_parameters": 17521921,
    "n_copies": 10,
    "info": "VGGNet L (17521921, 64, 14.040, 0.0189)",
    "info_short": "(VGGNet L, 64, 14.040, 0.0189)",
    "excess_negative": 100,
    "emit": false,
    "absorb": false,
    "loss": 0.01889071986079216
}
```

**Model groups**

Instead of saving single models, it is also possible to save multiple models and the interactions between them as a group.

In this case, a single file with extension .scnmgroup is created. This file is also an uncompressed .tar archive, and it contains one .scnm file for each model in the group, as well as a .json file that describes the interactions between the models.
